# Role of EXO1 nuclease activity in genome maintenance, the immune response and tumor suppression in *Exo1^D173A^* mice

**DOI:** 10.1101/2021.10.05.463200

**Authors:** Shanzhi Wang, Kyeryoung Lee, Stephen Gray, Yongwei Zhang, Catherine Tang, Rikke B. Morrish, Elena Tosti, Johanna van Oers, Paula E. Cohen, Thomas MacCarthy, Sergio Roa, Matthew Scharff, Winfried Edelmann, Richard Chahwan

## Abstract

DNA damage response pathways rely extensively on nuclease activity to process DNA intermediates. Exonuclease 1 (EXO1) is a pleiotropic evolutionary conserved DNA exonuclease involved in various DNA repair pathways, replication, antibody diversification, and meiosis. But, whether EXO1 facilitates these DNA metabolic processes through its enzymatic or scaffolding functions remains unclear. Here we dissect the contribution of EXO1 enzymatic versus scaffolding activity by comparing *Exo1^DA/DA^* mice expressing a proven nuclease-dead mutant form of EXO1 to entirely EXO1-deficient *Exo1^−/−^* and EXO1 wild type *Exo1^+/+^* mice. We show that *Exo1^DA/DA^* and *Exo1^−/−^* mice are compromised in canonical DNA repair processing, suggesting that the EXO1 enzymatic role is important for error-free DNA mismatch and double-strand break repair pathways. However, in non-canonical repair pathways, EXO1 appears to have a more nuanced function. Next-generation sequencing of heavy chain V region in B cells showed the mutation spectra of *Exo1^DA/DA^* mice to be intermediate between *Exo1^+/+^* and *Exo1^−/−^* mice, suggesting that both catalytic and scaffolding roles of EXO1 are important for somatic hypermutation. Similarly, while overall class switch recombination in *Exo1^DA/DA^* and *Exo1^−/−^* mice was comparably defective, switch-switch junction analysis suggests that EXO1 might fulfill an additional scaffolding function downstream of class switching. In contrast to *Exo1^−/−^* mice that are infertile, meiosis progressed normally in *Exo1^DA/DA^* and *Exo1^+/+^* cohorts, indicating that a structural but not the nuclease function of EXO1 is critical for meiosis. However, both *Exo1^DA/DA^* and *Exo1^−/−^* mice displayed similar mortality and cancer predisposition profiles. Taken together, these data demonstrate that EXO1 has both scaffolding and enzymatic functions in distinct DNA repair processes and suggest a more composite and intricate role for EXO1 in DNA metabolic processes and disease.

## INTRODUCTION

Exonuclease 1 (EXO1) is an evolutionarily conserved member of the XPG/Rad2 family of metallonucleases. EXO1 and its biochemical nuclease activity was initially identified in extracts of fission yeast undergoing meiosis and has since been shown to fulfill broader biological functions in mammalian cells during DNA damage responses (DDR), including DNA mismatch repair (MMR), DNA double-strand break repair (DSBR), and telomere maintenance (1–3). In immune B cells, EXO1 can also mediate non-canonical forms of MMR that promote antibody diversification through somatic hypermutation (SHM) and class switch recombination (CSR) (4). In germ cells, EXO1 mediates genetic diversity by promoting recombination during meiosis (5). Catalytically, EXO1 has 5’ to 3’ exonuclease and 5’-flap endonuclease activities. Yet, these seemingly distinct nuclease activities are now shown to be mechanistically integrated (6).

DNA mismatches are some of the most common forms of DNA damage, arising mainly during replication. MMR is the main pathway that removes misincorporated nucleotides that result from erroneous replication. During MMR, the coordination of mismatch recognition and excision is facilitated by MutS homolog (MSH) and MutL homolog (MLH) complexes. Briefly, MSH2 forms a complex with MSH6 (MutSα) to initiate the repair of single-base mispairs and insertion/deletions (IDLs) and with MSH3 (MutSβ) to initiate the repair of larger IDLs of two to four bases. Subsequent to mismatch recognition, MutSα and MutSβ interact with the MLH1-PMS2 (MutLα) complex in an ATP-dependent manner to initiate EXO1 mediated excision of the DNA strand carrying the mismatched base(s). The repair process is completed by DNA re-synthesis to fill the resulting single strand gap and ligation. Thus far, EXO1 is the only known exonuclease that facilitates the removal of the mismatch-carrying strand during MMR (7, 8). Through MMR, EXO1 improves replicative fidelity by over two orders of magnitude and reduces genomic mutator phenotypes (7, 9). During MMR, EXO1 directly interacts with MLH1, MSH2, and MSH3 both in yeast (10, 11) and mammalian cells (12, 13). As a result, it was speculated that EXO1 could fulfill both catalytic and structural roles during MMR but direct *in vivo* evidence for this idea remained elusive (14). Interestingly, a non-canonical error-prone subset of MMR occurs during B cell maturation. During this process, initiated by AID-induced U:G mismatches in germinal center B cells, EXO1 is believed to resect mismatches akin to its role during canonical MMR. However, when repairing AID induced mismatches in antibody V regions, this process subsequently diverges by the recruitment of error-prone polymerases, which were shown to be recruited by ubiquitinated PCNA (15, 16). Another repair mechanism that relies on EXO1 activity is DSBR, which repairs some of the most toxic forms of DNA damage. DNA end resection of DSBs in the 5’ to 3’ direction is a critical intermediate in this process. The ensuing single-stranded DNA (ssDNA) induces two inter-dependent responses, namely checkpoint signaling and initiation of homologous recombination (HR) mediated DNA repair. Studies in yeast and mammals (17, 18) suggest a two-step model for DSB processing. MRE11-RAD50-NBS1 (MRN) and CtIP initiate the ‘end-trimming’ of the DSB, which is then followed by the generation of longer stretches of ssDNA by either EXO1 or the BLM/DNA2 helicase-nuclease complex (17, 19). Two physiological and beneficial subsets of DSBR occur during B cell maturation through CSR and during meiosis in germ cells. Both processes are dependent on adequate upstream DSBR signaling, but they diverge considerably downstream. While meiosis relies on the error-free HR pathway for crossover formation, CSR relies on the error-prone pathway of non-homologous end joining (NHEJ) to generate intra-chromosomal switch region translocations and facilitate isotype switching. As a result of this divergence, the dependency of these two processes on EXO1-mediated resection remains unclear.

Deficiencies in MMR and DSBR associated processes can lead to chromosomal instability, infertility, neurodegeneration, tumorigenesis, and immune defects (20). Given its pleiotropic roles in the DNA damage response, it is expected that defects in EXO1 could also lead to some of these anomalies. Indeed, mice with an inactivating mutation in EXO1 had meiotic defects and sterility due to loss of chiasmata during metaphase (21), impaired SHM and CSR, loss of mismatch repair activity, and a 30 fold increase of mutation frequency in mouse ES cells (4–5). EXO1 inactivation also reduced overall survival in mice and caused an increased incidence of cancers, especially lymphomas (4–5). In humans, germline mutations in *EXO1* have been associated with hereditary nonpolyposis colorectal cancer/Lynch syndrome (HNPCC/LS) (23).

To dissect the scaffolding function from the enzymatic function of EXO1, a mouse line with the cancer-associated E109K mutation was previously generated and characterized (5). Residue E109 is a surface residue of the catalytic domain in the crystal structure (24) and, even though it does not directly affect the catalytic site, it had been reported to be essential for the nuclease activity when the N-terminal fragment of human EXO1 was examined biochemically (5, 25). *In vivo*, homozygous mutant *Exo1^EK/EK^* knock-in mice carrying the EXO1^E109K^ showed defects in the DNA damage response (DDR) and increased cancer predisposition. Unexpectedly the *Exo1^EK/EK^* mice showed normal activity in SHM, CSR, MMR, meiosis and were fertile, while *Exo1^−/−^* mice had defects in all of these processes. This was interpreted to mean that EXO1 had an essential scaffolding role, but its enzymatic activity could be replaced by some unknown factor during SHM, CSR, MMR, and meiosis (5). However, this conclusion was challenged by two subsequent independent biochemical observations, where for the first time full-length non-tagged EXO1^E109K^ was purified *in vitro*, and near WT activity was detected (26, 27). In contrast to the E109K mutation, these studies also confirmed that the EXO1^D173A^ mutation within the catalytic site, which had been previously reported to be nuclease-dead in yeast (28), also caused the clear inactivation of EXO1 nuclease activity *in vitro* (26, 27), offering a new alternative to reevaluate the nuclease and scaffold functions of EXO1 *in vivo*. Here, to separate the nuclease and scaffold functions of EXO1, a new knock-in mouse line carrying the EXO1^D173A^ mutation (*Exo1^DA^*) was generated, and the resulting phenotypes compared to *Exo1^−/−^* and *Exo1^+/+^* mice. Characterization of these animals confirms that EXO1 exhibits both scaffolding and catalytic functions, demonstrating that its exonuclease activity is essential in most DNA repair mechanisms (i.e. DDR, MMR, SHM and CSR) while it remains nonessential for meiosis, where EXO1 appears to play a critical structural role.

## MATERIALS AND METHODS

### Mice and MEF strains

The generation of Exo1^−/−^ mice has been described previously (4). *Exo1^DA/DA^* mice were generated using CRISPR/Cas9 technology (58). In brief, the mixture of guide RNA targeting to the adjacent area of Exo1 D173 coding region on exon6 (The CRISPR targeting sequence is gactctgacctcctcgcattTGG), Cas9 mRNA, and donor DNA carrying the approximately 1kb homologous arms at each side and the surrounded desired DNA mutation were injected into zygotes of C57BL/6J mice and then the injected fertilized eggs were transferred into pseudopregnant CD1 female mice for producing pups. The resulting offspring were genotyped by PCR and Sanger sequencing to identify the founders carrying the desired Exo1 D173A mutation. The founders (F0) were mated with wild-type *C57BL/6J* mice, and then the resulting germline transmitted F1 heterozygotes were bred to establish study cohorts. When indicated, *Exo1^+/+^*, *Exo1^+/DA^*, and *Exo1^+/−^* mice were further crossed to *Big Blue* transgenic mice (58) or *p53+/−* mice (5). Mouse embryonic fibroblasts (MEFs) were isolated from embryos of pregnant mice at day 13.5 post-conception. MEF cell lines were passaged 4 – 6 times and cryopreserved. All protocols were performed in accordance with the Animal Care and Use Committee of Albert Einstein College of Medicine (AECOM) under the direction of Public Health Service (PHS) Policy on Humane Care and Use of Laboratory Animals.

### Analysis of *cII* mutation frequencies

*Exo1^+/+^*, *Exo1^+/−^* and *Exo1^+/DA^* mice were crossed to Big Blue transgenic mice (56) to generate *Exo1^+/+^/BigBlue*, *Exo1^−/−^/BigBlue*, and *Exo1^DA/DA^/BigBlue* mice. Mutations in genomic DNA from the spleen and liver were detected using the λ Select-cII Mutation Detection System for Big Blue Rodents (Stratagene) as we have done in the past (58).

### Metaphase Analysis

MEFs were collected after 4 hours of treatment with 10 ng/mL colcemid and treated with 75 mM KCl for 20 min. After centrifugal collection, MEFs were fixed with a 3:1 (v/v) mixture of methanol and acetic acid. Fixed cells were then transferred onto a microscope slide and mounted with VECTASHIELD Mounting Media with DAPI (Vector Labs). Subsequently, cells were analyzed using a fluorescent microscope.

### CPT Treatment and Immunofluorescence

Primary MEFs were grown on coverslips and treated with 1 μM CPT or DMSO (control). After 1 h, the drug was removed and cells were pre-extracted for 5 min on ice in 10 mM Pipes buffer (pH 6.8) containing 300 mM sucrose, 50 mM NaCl, 3 mM EDTA, 0.5% Triton X-100, and Protease Inhibitor Mixture (EDTA-free; Roche) before fixation in 2% (wt/vol) paraformaldehyde for 15 min at 25 °C. After fixation, cells were washed with PBS and then were blocked with 5% (wt/vol) BSA and 0.1% Triton X-100 in PBS before staining with mouse anti-gH2AX (Cell Signaling), rabbit anti-RPA pS4/S8 (Bethyl), and DAPI (Vector) for 1 h. After washes in PBS + 0.1% Triton X-100, Alexa 488 goat anti-mouse/rabbit, and Alexa 598 goat anti-mouse/rabbit (Molecular Probes) were used as secondary antibodies. Images were acquired using a Bio-Rad Radiance 2100 (Nikon Eclipse E800) microscope using Lasersharp 2000 software (Zeiss).

### MNNG Treatment

After crossing all models presented here to *p53^+/−^* mice, MEF cell lines were generated and pretreated with 20 μM O6-Benzylguanine (O6BG, Sigma) before the addition of MNNG to inhibit the repair of O6meG adducts by O-6-methylguanine-DNA methyltransferase (MGMT, Sigma). After 48h MNNG treatment, relative cell viability was determined using Thiazolyl Blue Tetrazolium Bromide (MTT)-conversion (Sigma) as per the manufacturer’s instructions. The absorbance of converted dye was measured at a wavelength of 570 nm using a Perkin Elmer Victor X5 plate reader. Cell viability was calculated relative to DMSO-treated cells incubated in parallel.

### Analysis of Murine Tumors and Survival

Mice were routinely observed until they became moribund and/or signs of tumour development occurred and required euthanasia. The Kaplan-Meier method was used to compare the overall survival curves of the mice, using the log-rank analysis to evaluate significance. All tumors were processed for paraffin embedding, and sections were prepared for H&E staining and evaluated by a pathologist. Statistical analysis of tumour incidence was performed using the Fisher’s exact test.

### Immunization and Hypermutation Analysis on V_H_186.2

Age-matched mice (*Exo1^+/+^*, *Exo1^−/−^*, and *Exo1^DA/DA^*) were immunized intraperitoneally with 200 μg NP(33)-CGG (BioSearch Technologies) on alum. For primary response analysis, animals were sacrificed, and spleens were removed 10 days after a single immunization. Subsequently, RNA was extracted using Purelink RNA Mini Kit (ThermoFisher) and cDNA was generated using AccuScript High Fidelity 1st Strand cDNA Synthesis Kit (Stratagene). The *Mus musculus* IGHV1-72*01 allele (IMGT accession number J00530:206-499), referred here as V_H_186.2, was amplified using a nested PCR as previously described (16), and PCR products were gel purified and stored in 50 μL buffer containing 10mM Tris (pH 8.0) until submission to TruSeq^®^ DNA library preparation (Illumina) following manufacturer’s instructions. After quality control by Bioanalyzer (Bio-Rad), libraries were pair-end sequenced on a MiSeq Sequencer (Illumina) at Einstein Epigenetics Facility. Using an in-house computational pipeline based on Kernel written in ultra-fast C++ and a GUI written in Java, Illumina pair-reads were assembled and collapsed according to their unique molecular identifier (UMI) to reconstruct unique overlap VH186.2DJ regions. These sequences in FASTA format were then submitted to IMGT/HighV-QUEST (61) for VH186.2-D-J usage and CDR3 clonality identification. Then, customized Python and R scripts were used for mutational analyses after randomly resampling during 1000 iterations one single sequence per CDR3 clone, allowing us to increase our sensitivity for unique mutations, which are expected to derive from independent mutational events, and avoid overestimating mutation frequencies by repeated counting of non-unique mutations inherited within the clonal CDR3 lineage. For each group of *Exo1^+/+^*, *Exo1^−/−^*, and *Exo1^DA/DA^* iterations, frequencies of mutation were normalized to the potential maximum number of the corresponding mutation subtype as we have previously described (62), which corrects for sequence composition and size of CDR3 repertoire.

### *Ex Vivo* assay of class switching

Splenic cells were isolated from immunized animals (*Exo1^+/+^*, *Exo1^−/−^*, and *Exo1^DA/DA^*). Red blood cells and T cells were removed by ammonium chloride and complement-dependent lysis with details described as previously (16). The rest of the cells were stimulated either using 50 μg/ml of LPS from Sigma-Aldrich (St. Louis, MO) or using 50 μg/ml of LPS plus 50 ng/ml of mouse IL-4 from R&D Systems (Minneapolis, MN). After 4 days in complete RPMI media at 37°C, cells were stained against surface IgM (ImmunoResearch) plus IgG1 (SouthernBiotech) or IgM plus IgG3 (SouthernBiotech), which was followed by FACS analysis.

### Switch Junction Analysis

After LPS stimulation for 4 days, splenic B cells were collected for genomic DNA isolation using the DNeasy kit (Qiagen). Sμ–Sγ3 regions were amplified using nested PCR with details and primers as previously described (63). PCR products were purified with QIAquick PCR purification kit (Qiagen), ligated to Topo Blunt vector (Invitrogen) and transformed in *E. coli* chemical competent cells. Plasmids from individual *E. coli* colonies were purified and sequenced for the inserted switch junctional regions. Switch junctions were analyzed by alignment using BLAST2seq with the automatic low-complexity filter disabled and using consensus sequences from the NC_000078.6 GenBank record.

### Meiosis

Analysis of Meiotic Prophase I was performed as previously described (64, 65). After Chromosome spreading, the slides were washed, blocked, and then incubated with a primary antibody against synaptonemal complex protein 3. After subsequent washing and blocking, the slides were incubated with a secondary antibody conjugated to fluorescein. After further washing, slides were analyzed using an upright fluorescent microscope (Olympus BX61 with a CoolSNAP HQ camera).

## RESULTS

### Generation of nuclease-dead *Exo1^D173A^* mice

To create an inactivating nuclease-dead EXO1 mutation, we generated a novel knock-in mouse line that carries the EXO1^D173A^ mutation (termed *Exo1^DA^*) by CRISPR/Cas9 technology in fertilized oocytes isolated from C57BL/6 female mice (Figure S1A). The *Exo1^DA^* mutation was verified in the offspring by PCR and Sanger sequencing (Figure S1B). This mutation was previously shown to have no detectable nuclease activity biochemically (26, 27). Notably, the EXO1^D173^ residue is evolutionarily conserved in both fission and budding yeast as well as mouse and humans (Figure 1A, 1B). Furthermore, structural modeling of the resolved EXO1 crystal structure shows that residue EXO1^D173^ is embedded in the exonuclease active site and is in direct contact with DNA moieties (6), which predicts a pronounced anomaly in the EXO1 nuclease active site as a consequence of the EXO1^D173A^ mutation (Figure 1C). However, such a perturbation is not expected to affect protein folding and stability, DNA binding, or MMR protein interactions as demonstrated by the distinct domain dependencies (Figure 1B-C) and this is supported by studies in yeast and mammalian cells (26, 27, 29–31). Consistent with this, we observed stable expression of mutant Exo1^DA^ protein comparable to wild type EXO1 by Western-blot analysis, compared to complete protein loss in previously characterized *Exo1^−/−^* mice (5) (Figure S1C).

**Figure 1:**
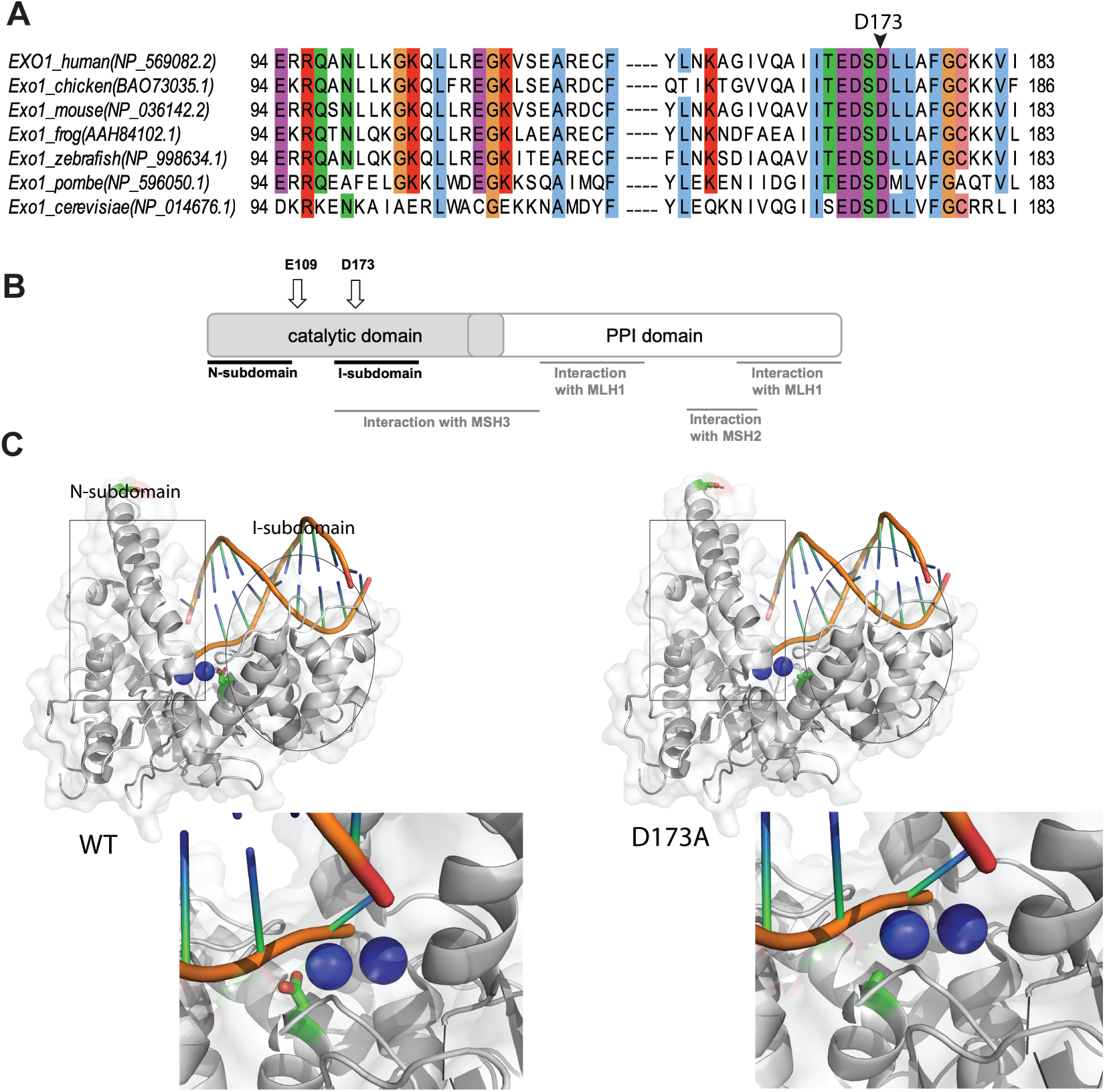
Modeling EXO1 mutations in mice. A) Jalview ClustalW alignment of EXO1 from the denoted species shows high evolutionary conservation of the D173 residue. B) Schematic structure depicting human EXO1 domains and binding partners MSH3, MSH2 and MLH1 which help mediate some of EXO1 non-catalytic scaffolding functions. C) Crystal structure of the N-terminal nuclease (enzymatic) domain (dark grey) of human EXO1 (left), with the inset showing a 180° rotation of the active site; and D173A (right), with the inset showing a 180° rotation of the active site. Active site metals shown in cyan. (PDB: 3QEB). The insets highlights the difference between the proximity of the metal ions (2 cyan circles) in the active site to the acidic D173 amino acid residue (green and red) compared to the conformational change generated by the neutral A173 amino acid residue (green) mutation.

### The catalytic activity of EXO1 is predominant over scaffolding functions in canonical DDR

To measure the integrity of canonical error-free MMR function *in vivo*, *Exo1^+/+^*, *Exo1^DA/DA^*, and *Exo1^−/−^* mice were crossed to *BigBlue* mice (5, 32). The accumulation of mutations in the *cII* reporter gene was analyzed in genomic DNA from the liver and spleen in all three cohorts. We observed that mutation rates were consistently 2-3 times higher in both *Exo1^−/−^* and *Exo1^DA/DA^* mutant mice compared to *Exo1^+/+^* mice, thereby confirming that the catalytic role of EXO1 is essential for canonical error-free DNA mismatch repair in mammalian cells *in vivo* (Figure 2A).

**Figure 2:**
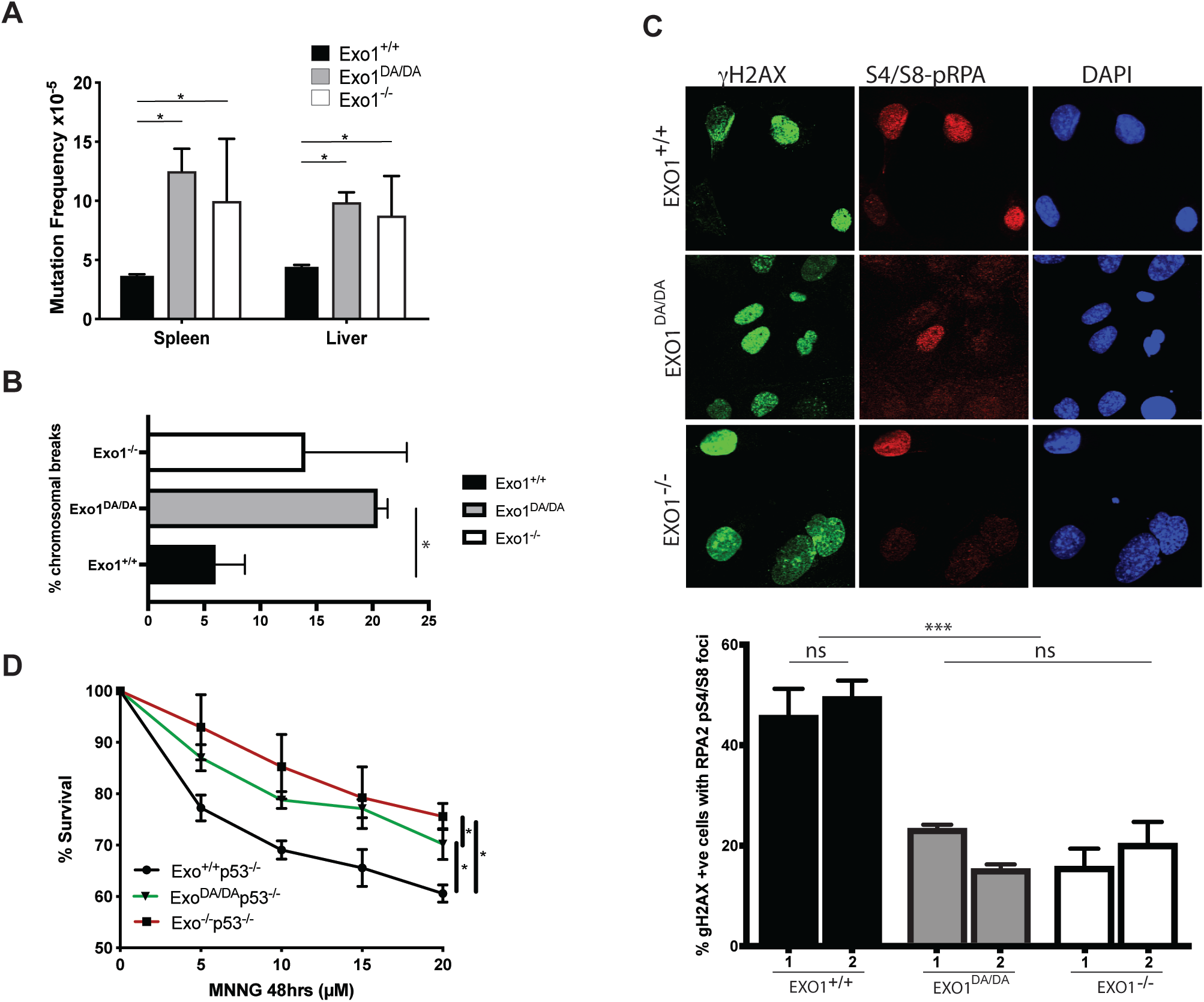
*in vivo* nuclease activity of EXO1 is required for canonical DDR. A) *cll* reporter gene mutation frequencies in spleen and liver of *Exo1^+/+^*, *Exo1^DA/DA^*, and *Exo1^−/−^* mice (mice/genotype). Mutation frequency is increased by two-to threefold in *Exo1^−/−^* and *Exo1^DA/DA^*. B) Quantification of the number of chromomal breaks from metaphase spreads of the denoted *Exo1* genotypes. At least two independent cell lines we used per genotype and more than 100 cells where counted per cell line. Significance was established using T-test (*P < 0.01). C) Representative photographs showing γH2AX, pS4/S8-RPA2, and DAPI stained MEFs after camptothecin treatment (2 h, 1 μM) of the indicated genotypes. (Mag: 1,000×; >80 cells were visualized per genotype). Histogram of the rates of γH2AX-positive (DSB marker) cells with RPA2-pS4/S8 foci (ssDNA marker) in MEFs of the indicated genotypes. ns, not significant; ***P < 0.001. D) Cellular survival curves of MEFs from *Exo1^+/+^*, *Exo1^DA/DA^*, and *Exo1^−/−^* mice in a p53-null background following 48h treatment with the denoted concentration of MNNG (x-axis). Error bars represent ± SD; n.s., not significant; *P-value < 0.01.

In order to investigate the role of EXO1 in DSBR, we established primary mouse embryoninc fibroblasts (MEFs) from *Exo1^+/+^*, *Exo1^DA/DA^*, and *Exo1^−/−^* mice and quantified chromomal breaks in metaphase spreads (Figure 2B). Compared to control *Exo1^+/+^*, both *Exo1^−/−^* and *Exo1^DA/DA^* MEFs exhibited a similar increase in the percentage of cells carrying chromosomal breaks, suggesting that EXO1 nuclease is also essential for DSBR and chromosomal stability. To examine whether the accumulation of these DSBs responded to defects in EXO1 functions during end resection to initiate DSBR, we exposed *Exo1^+/+^*, *Exo1^DA/DA^*, and *Exo1^−/−^* primary MEFs to camptothecin (CPT), which induces DSBs specifically in S-Phase as a result of increased replication fork collapse intermediates. We then investigated the frequency and co-localization in foci of hyperphosphorylated RPA (S4/S8-pRPA) and γH2AX, which are targeted by DNA damage-activated protein kinases of the PI3-family like ATM/ATR as a reaction at DSBs (Figure 2C). This allowed us to score DNA end resection events during DSB repair, where both *Exo1^−/−^* and *Exo1^DA/DA^* MEFs showed significantly reduced co-localization of activated pRPA-S4/S8 and γH2AX (Figure 2C). These results indicate impaired DSB-resection in response to CPT treatment and suggest a critical role for the catalytic activity of EXO1 during DSBR in mitotic cells.

MMR proteins also play a critical role in mediating adequate DDR signaling following genotoxic stress either directly by acting as DNA damage sensors to activate DDR signaling networks (direct signaling model) or indirectly by sequentially escalating a mutation in a ssDNA intermediate to make a dsDNA break, which in turn triggers cell cycle arrest and cell death (futile cycle model) (33). To investigate whether the EXO1^D173A^ mutation could impair the eliciting of a G2-phase checkpoint arrest and apoptosis in response to S_N_1 DNA methylators such as MNNG (34, 35), we compared the survival of immortalized MEFs on a *p53^−/−^* background isolated from *Exo1^+/+^*/*p53^−/−^*, *Exo1^DA/DA^*/*p53^−/−^* and *Exo1^−/−^*/*p53^−/−^* mice. We found that *Exo1^−/−^* and also *Exo1^DA/DA^* p53-deficient MEFs, although slightly less robustly in the *Exo1^DA/DA^* mutants, displayed increased resistance to MNNG treatment compared to *Exo1^+/+^* MEFs, suggesting inefficient DDR signaling in both EXO1 mutant backgrounds (Figure 2D). In conjunction with previous mechanistic studies showing the emergence of ssDNA and dsDNA breaks following MNNG exposure (30–32), these results indicate that the nuclease activity of EXO1 facilitates MMR-mediated DDR signaling.

### EXO1 catalytic function is important for survival and tumor suppression

DNA mismatches and DSBs are considered the most abundant and harmful of all genotoxic lesions, respectively. Failure to adequately repair these lesions can have severe adverse effects on organisms, including genomic instability and induction of tumorigenesis (36–38). To study the long-term effects of EXO1 nuclease inactivation on survival and cancer susceptibility, *Exo1^+/+^*, *Exo1^−/−^* and *Exo1^DA/DA^* mouse cohorts were monitored for a period of up to 26 months. While *Exo1^+/+^* mice exhibited the expected overall survival with 50% of mice still alive at 24 months of age, both *Exo1^−/−^* and *Exo1^DA/DA^* mice had a comparable decreased survival rate with 50% of mice dead at 16 months of age (Figure 3A). These accelerated death rates in *Exo1^−/−^* and *Exo1^DA/DA^* mice were due to increased cancer predisposition at a comparable incidence (Figure 3B). The majority of *Exo1^DA/DA^* mice died of lymphoma (58%, n=14/24) between six and 22 months of age (Figure 3C, D). In addition, two mice developed gastrointestinal adenomas (8%, n=2/24) and three mice developed liver carcinoma (10%, n=3/24). In two mice, both lymphoma and sarcoma or lymphoma and fibroma were found. Five mice died before an autopsy could be performed (21%, n=5/24). The tumor spectrum on *Exo1^−/−^* was largely identical to that of *Exo1^DA/DA^* mice (Figure 3C,D). In addition, similar to *Exo1^−/−^*/*p53^−/−^* mice, the introduction of a p53 knockout allele led to the rapid deaths of *Exo1^DA/DA^*/*p53^−/−^* mice between two to three months of age due to the development of T cell lymphoma (100%, n=17/17) ((5) and not shown). Taken together, these results indicate that an intact nuclease activity is essential for EXO1 tumor suppressor functions.

**Figure 3:**
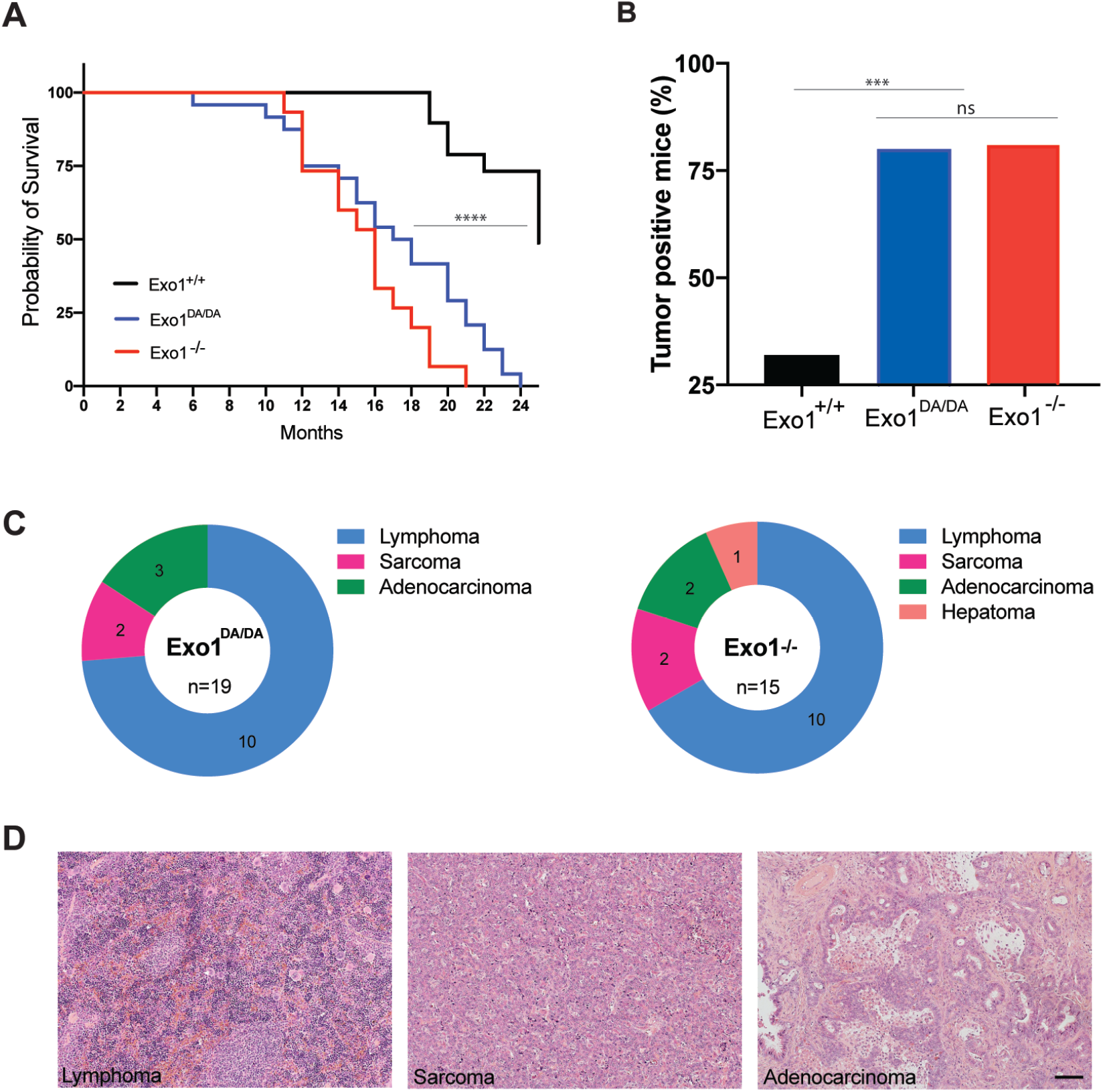
The role of EXO1 nuclease-dead mutant on tumorigenesis. A) Animal survival curve of the denoted genotypes showing faster death of *Exo1^DA/DA^* (n=24) and *Exo1^−/−^* (n=15) mice compared to WT (n=47). B) *Exo1^−/−^* (n=15) and *Exo1^DA/DA^* (n=19), which, as shown in B), had a considerably higher tumor positive incidence compared to WT (n=20). C) representative distribution of the tumor profiles of the samples quantified in B). D) representative H&E stains of the tumor samples denoted in C). Bar is 50 μm.

### Catalytic and structural functions of EXO1 are required for non-canonical DDR during antibody diverfication

To delineate the precise contribution of EXO1 structural and/or catalytic functions during SHM, we examined error-prone repair of the VH186.2 heavy chain variable region during primary immune responses in the *Exo1^+/+^* (n=7), *Exo1^DA/DA^* (n=8), and *Exo1^−/−^* (n=3) cohorts (Figure 4A and B). In the absence of either the EXO1 protein or its nuclease activity, a significant imbalance of mutations at G:C and A:T pairs was revealed when frequencies of mutation were compared to those normally observed in *Exo1^+/+^* mice. While mutations at A:T pairs and error-prone Polη hotspots were significantly reduced, the remaining mutations were not only restricted but also increased at G:C pairs and AID hotspots in both *Exo1^−/−^* and *Exo1^DA/DA^* mice (Figure 4A). This pattern of nucleotide substitutions strongly suggests that G:C mutations are due to replication over the initial AID-induced U:G lesion without proper EXO1-dependent resection of the surrounding area and, therefore, without the consequent induction of A:T mutations by the recruitment of error-prone DNA polymerases like Polη. Interestingly, the latter defect in error-prone repair at A:T sites was less dramatic in the presence of the mutant Exo1^DA^ protein than in its complete absence (Figure 4A), suggesting that some intermediate scaffolding functions might still persist in *Exo1^DA/DA^* B cells. Consistently, while a reduction in the spread of A:T mutations across the length of the VH186.2 region, specially at Polη hotspots, could be observed in both EXO1-compromised models, *Exo1^−/−^* mice showed a more clear altered phenotype with its few remaining A:T mutations distributed around CDR2 (Figure 4B). On the other hand, the overall distribution and preference for mutations at G:C sites in the CDRs was maintained and even increased at non-hotspot G:C sites in *Exo1^−/−^* and *Exo1^+/+^* mice (Figure 4B). Overall, analysis of *in vivo* SHM at immunoglobulin genes suggests that the structural and enzymatic functions of Exo1 work in tandem to provide adequate error-prone MMR around AID-induced U:G mismatches.

**Figure 4:**
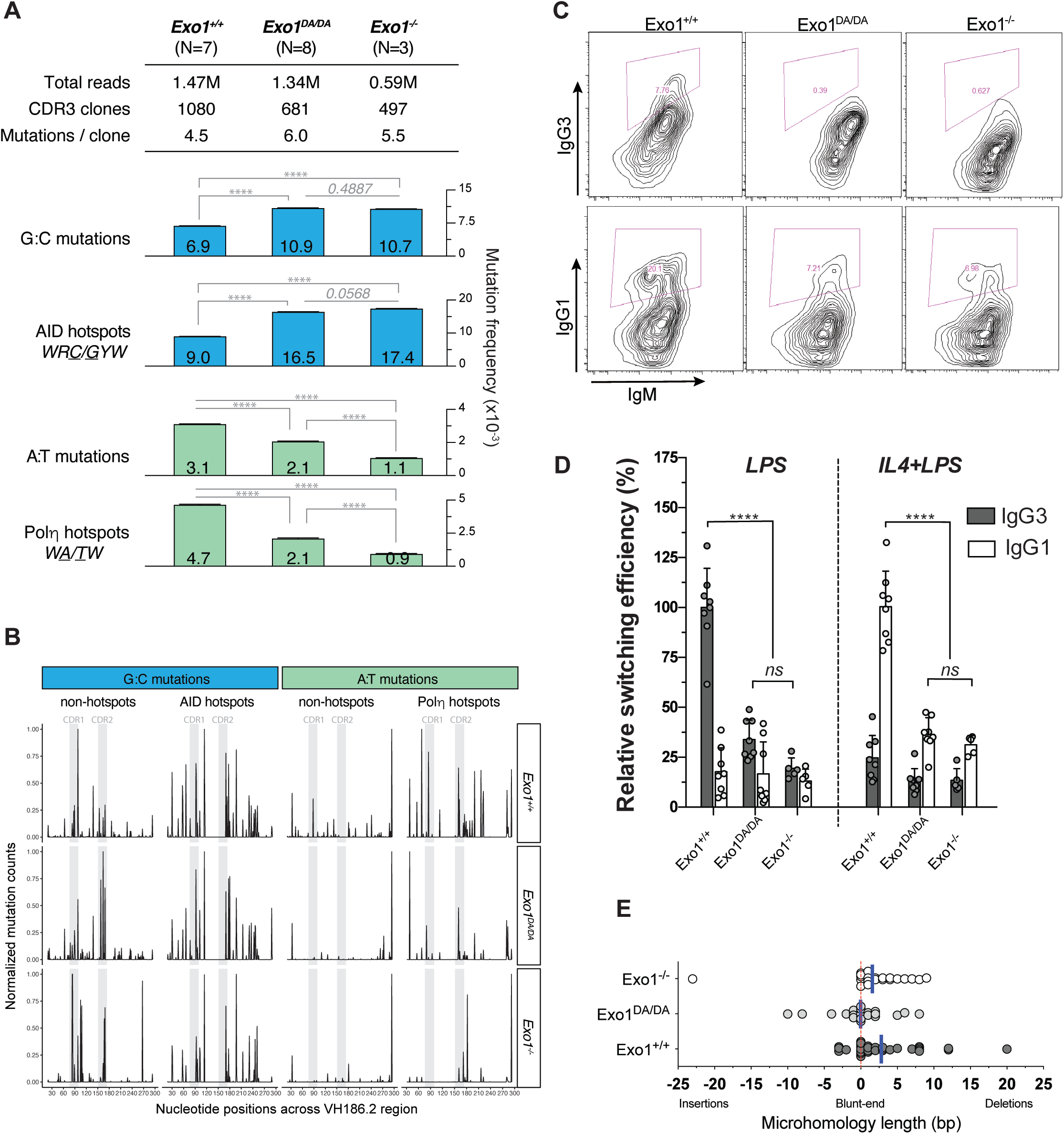
Abrogation of EXO1 or its nuclease activity impairs antibody diversification. A) Global analysis of *in vivo* SHM at the VH186.2 region during primary response of NP-immunized mice. For comparison of average mutation frequencies of *Exo1^−/−^* and *Exo1^DA/DA^* to WT, contingency tables were assigned to the mutation data and Pearson’s Chi-squared test with Yates’ continuity correction was applied. B) Distribution of unique mutations across the VH186.2 region and its CDRs as observed during the 1000 iterations that were performed for each genotype group and normalized within each mutation subtype to maximum of 1. Mutations in G:C or A:T residues are split and are colored according to the hotspot definition for AID (WR<u>C</u>/<u>G</u>YW) or poln (W<u>A</u>/<u>T</u>W). C) Representative flow cytometry profiles analyzed by FloJo showing CSR efficiency in the three denoted mouse cohorts. D) Summary of *ex vivo* relative CSR efficiencies in WT (n=8), *Exo1^DA/DA^* (n=8) and *Exo1^−/−^* (n=5) mice. As reference, mean efficiency of switching in the WT group to IgG3 after stimulation with LPS or to IgG1 after stimulation with IL4+LPS were defined as 100% in two independent experiments. Relative percentages of switching to each isotype by animal and stimulation cocktail are depicted by dots, and bars represent mean ± SD of the replicates. The background switching to IgG1 after LPS stimulation and to IgG3 after IL4+LPS stimulation are also shown. E) Distribution of blunt breakpoints (0 bp), microhomologies (from 1 to >10 bp in length) or insertions (represented as negative integers) in Sμ-Sγ3 switch junctions cloned and sequenced in *ex vivo* LPS-stimulated splenic B cells from the different genotype groups (n>22 per group). Dots represent individual Sμ-Sγ3 switch junctions, and blue bars denote mean length in the distribution of blunt junctions, microhomologies and insertions within each group.

In B lymphocytes, generating and resolving DSB intermediates are critical to ensure diversity of antibody effector functions through CSR (38–40). To determine whether the aforementioned lack of adequate ssDNA generation observed in EXO1 nuclease-deficient cells (Figure 2) bears any physiological cellular phenotype in B cells, we conducted *ex vivo* CSR experiments where DSBs at Ig switch regions are obligate intermediates in the isotype recombination process (38). To this end, B cell splenocytes were isolated from *Exo1^+/+^* (n=8), *Exo1^−/−^* (n=5), and *Exo1^DA/DA^* (n=8) mice and subjected to lipopolysaccharide (LPS) to induce IgM to IgG3 isotype switching or LPS and interleukin-4 (IL-4) to induce IgM to IgG1 isotype switching ex vivo (Figure 4C). Notably, both *Exo1* mutant cohorts exhibited a significant decrease in CSR efficacy towards both IgG3 and IgG1, compared to *Exo1^+/+^* cohorts (Figure 4C and D), suggesting that EXO1 nuclease activity is critical for CSR. Next, to further explore whether EXO1 nuclease was required not only to generate the switch region DSBs themselves but also to resolve them, we cloned and sequenced Sμ-Sγ3 junctions from *Exo1^+/+^* and *Exo1^−/−^* and *Exo1^DA/DA^* mutant splenic B cells stimulated with LPS plus IL-4 for 72 hours. Switch region DSBs are known to be primarily resolved by classical NHEJ (cNHEJ), though a significant subset is also repaired through microhomology-mediated alternative-NHEJ (alt-EJ), where between 1 to 30 base pairs of ssDNA microhomologies flanking the AID-induced DSB guide the aNHEJ repair process (41, 42). We could observe that Sμ-Sγ3 junctions in *Exo1^DA/DA^* cells were different from those in *Exo1^+/+^* or *Exo1^−/−^* cells (Figure 4E). While *Exo1^+/+^* cells showed the expected range of 2-20bp microhomology at Sμ-Sγ3 junctions, *Exo1^−/−^* and *Exo1^DA/DA^* cells exhibited a slight decrease in long microhomologies. Furthermore, *Exo1^DA/DA^* cells were further skewed towards the use of blunt joints, with a clear increase in the number and length of insertions compared to the other two groups (Figure 4E). This resembles the phenotype previously observed with the depletion of the DNA end-processing factor CtIP (43), although this notion has now been contested (44). Taken together, these results appear to suggest that while EXO1 nuclease activity contributes to the end resection mechanisms that promote microhomology-mediated aNHEJ during CSR, a catalytically-dead protein may interfere with other factors that otherwise might participate and partially compensate during aNHEJ in the complete absence of EXO1.

### The structural function of EXO1, but not its catalytic activity, is critical for meiosis

Like CSR, meiosis is another example of beneficial DSBs in normal physiology. We had previously shown that *Exo1^−/−^* mice of both sexes were sterile (5). Interestingly, *Exo1^DA/DA^* male and female mice are fertile, suggesting normal meiotic progression. To observe crossover events in pachynema, we employed two well-characterized markers of these sites: MLH1 and CDK2 (45–47), which colocalize on the paired chromosomes and the telomeres during meiotic crossing over of mouse cells. In all our genotypic cohorts, MLH1 and CDK2 localized to crossover sites and CDK2 additionally localized to telomeres (Figure 5). These crossover markers remained associated with SYCP3, chromosome cores, and/or telomeres at a frequency and intensity that is indistinguishable between *Exo1^+/+^* and *Exo1^DA/+^* and *Exo1^DA/DA^* mice (Figure 5), indicating normal meiosis progression through prophase-I. In addition, by analyzing and quantifying metaphase spreads, we could show that the number of chiasmata and bivalent chromosomes was also comparable in all groups (Figure 5). This is in contrast to our previous findings in *Exo1^−/−^* mice in which metaphase spreads predominantly displayed abnormal spindle structures with mostly achiasmatic univalent chromosomes (5). Altogether, these results suggest that EXO1 nuclease activity, but not its structural function, is dispensable during meiotic progression.

**Figure 5:**
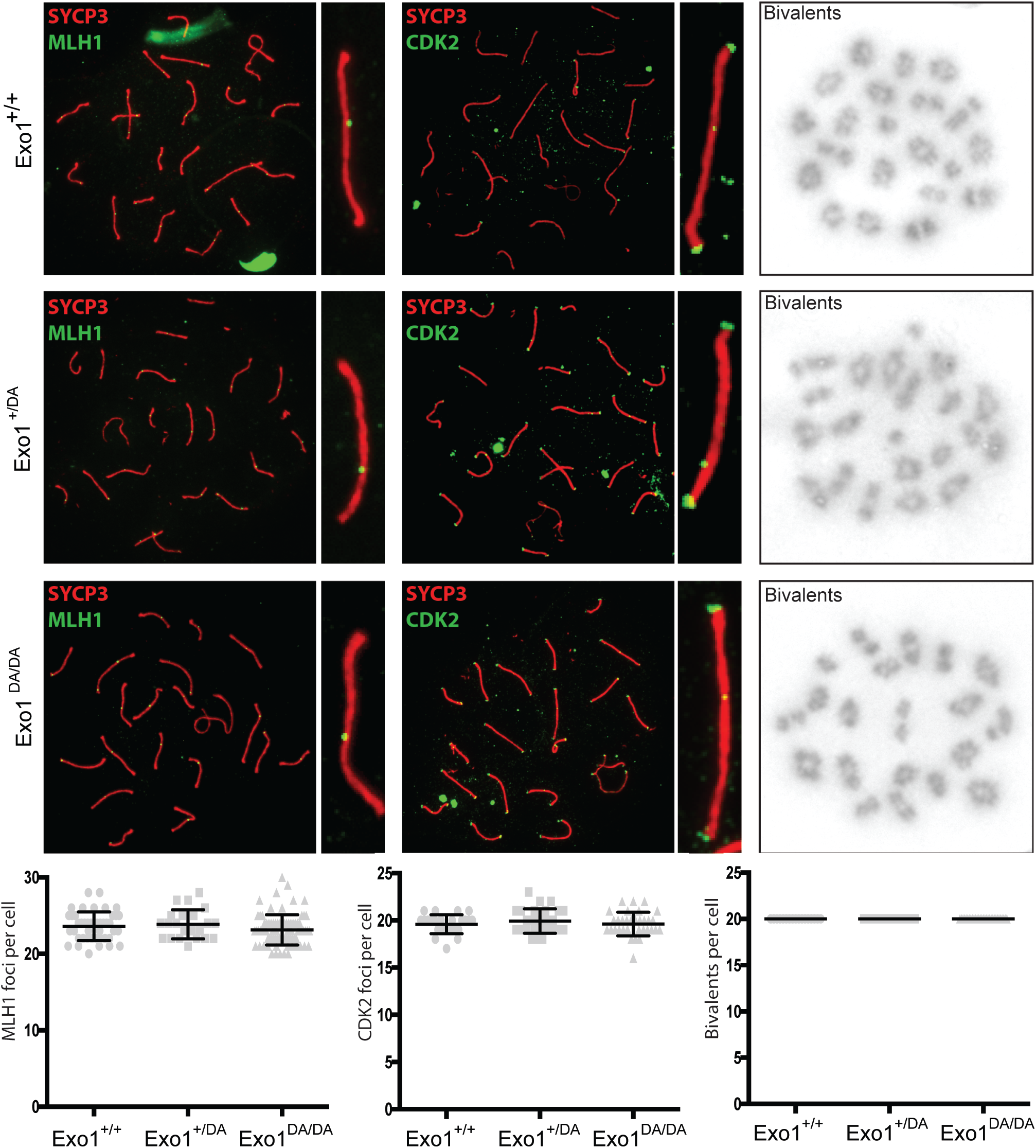
EXO1 nuclease activity is dispensable for meiosis during spermatogenesis. MLH1 and CDK2 (green) localize to the synaptonemal complex SYCP3 (red) in WT (top), *Exo1^DA/+^* (middle), and *Exo1^DA/DA^* (bottom) in pachytene spermatocytes. Left-hand panels show merged red and green channels, while the right-hand inset shows representative merged channels of a sample chromosome showing MLH1 and CDK2 at crossover sites and additional telomere-associated CDK2. Most right panels show the quantification of bivalent chromosome pairing during pachytene. The bottom charts shows the quantitation of foci for each genotype at pachynema. Counts for WT, *Exo1^DA^/+*, and *Exo1^DA/DA^* are not statistically different from each other (by unpaired t-test with Welch’s correction). Values given are the number of foci per nucleus ± s.d.

## DISCUSSION

As exemplified in this study, EXO1 fulfills many complex even contradictory functions ranging from maintaining genomic stability to promoting immune diversity as well as meiotic recombination. We specifically were interested in dissecting the distinct contributions of the catalytic and scaffolding functions of EXO1 in each of these genome maintenance processes *in vivo* by comparing the phenotypes of *Exo1^+/+^*, *Exo1^DA/DA^*, and *Exo1^−/−^* mice. The N-terminal fragment of EXO1 encodes the nuclease domain bearing the HNPCC-associated E109K mutation, which apparently compromised EXO1 nuclease activity *in vitro* when assayed using tagged N-terminal fragments of both mouse and human EXO1 (5, 25). However, recent studies in which the authors were finally able to isolate the full-length human EXO1^E109K^, without using any N-terminal tag, showed retention of significant nuclease activity (26, 27). A possible explanation for these conflicting observations could be related to structural instability in the EXO1^E109K^ mutant N-terminal fragment, together with the presence of a 6xHis-tag used for purification, which could both interfere with EXO1 enzymatic activity. Consistent with these observations, canonical and non-canonical MMR (SHM and CSR in B cells) were not affected in *Exo1^EK/EK^* mice. Yet, *Exo1^EK/EK^* mice showed increased cancer predisposition and displayed defective DDR and DSBR (5). This unexpected limited phenotype might be related to the recently indentified overlap of the EXO1 nuclease domain with a PAR-interaction (PIN) domain, which has been proposed to facilitate recruitment and resection functions of EXO1 during DSBR (48). Here, we bypassed those issues by creating and characterizing a novel *Exo1^DA/DA^* knock-in mouse model carrying the nuclease-dead D173A mutation that represented a definitive *in vivo* system allowing us to more precisely dissect potential structural and catalytic functions of EXO1 in different mammalian genome maintenance processes.

Our results suggest that EXO1 may have a 3-tier function in the various DDR processes in which it participates. This 3-tier approach depends on differential requirements for the catalytic vs. scaffolding domains of EXO1: catalytic, scaffolding, or both. They also seem to suggest that the complexity of the DDR mechanism somehow directs what activities of EXO1 are needed. In tier-1, we show here that the role of EXO1 in canonical DDR pathways, such as error-free MMR and DSBR, are primarily mediated by EXO1 nuclease activity. In our assay measuring the MMR of spontaneous genomic mutations, mutation frequencies were similar between EXO1 depleted (*Exo1^−/−^*) and EXO1 nuclease-dead (*Exo1^DA/DA^*) mouse models. Similarly, DSBR analyses through DNA end resection assays showed indistinguishable responses in these same mouse models. Furthermore, *Exo1^DA/DA^* and *Exo1^−/−^* mice exhibited similarly reduced survival due to the development of lymphomas, sarcomas, and gastrointestinal adenocarcinomas. These data suggest that error-free DNA repair through MMR and DSBR requires EXO1 resection activity and little or no distinct effects from its scaffolding function. Or that these two activities are completely epistatic or redundant in these particular repair mechanisms.

That EXO1 is involved in two dichotomous DDR pathways, namely error-free and error-prone MMR and DSBR, has been well documented (49, 50). SHM and CSR are two immune diversification mechanisms in B cells that specifically require non-canonical error-prone MMR and DSBR, respectively (51, 52). What was unexpected, though, is that during these two error-prone DNA repair processes, EXO1 seems to depend on both its catalytic and non-catalytic scaffolding functions (tier-2). EXO1 seems to initiate both SHM and CSR processes through its nuclease activity in the first instance. That is why our SHM analysis revealed a general increase in replication-fixed unrepaired C:G mutations and a decrease in error-prone introduced A:T mutations in both *Exo1^DA/DA^* and *Exo1^−/−^* models, which is in line with other MMR factor deletions, including MSH2 or MSH6 depletion (51, 52). Likewise, the overall decrease of IgM to IgG1 and IgG3 switching was comparable in the two mutant Exo1 mouse models, suggesting that the overall instigation of DSBs at switch junctions and their resolution by cNHEJ highly depend on EXO1 nuclease activity. However, upon further analysis of SHM mutation patterns, we could observe that while the increase in G:C mutations fixed by replication at AID-associated hotspot mutations were comparable between *Exo1^DA/DA^* and *Exo1^−/−^*, the recruitment of error-prone mechanisms and Polη-mediated mutations at A:T sites where only intermediary impaired in *Exo1^DA/DA^* compared to *Exo1^−/−^* B cells. These data suggest that during complex error-prone MMR repair, additional scaffolding functions of EXO1 are also needed. The rationale for how that could happen include: i) a passive reason, where nuclease-dead EXO1 could be recruited to sites of DNA mismatches, but its lack of nuclease activity could act as a dominant negative, thereby preventing subsequent downstream MMR signaling. However, this rationale should also apply for canonical MMR as well, which we did not observe in our assays. Another explanation could be the result of ii) an active scaffolding function of EXO1, which diversifies its function. In this sense, PCNA was shown to bind and increase EXO1 processivity at DSB sites (53), and PCNA is required for A:T mutagenesis during error-prone MMR through its recruitment of Polη (16). It is plausible that PCNA might still be recruited by the retained scaffolding functions of the mutant Exo1^DA^ protein and result in partial functionality, which would not otherwise occur in the complete absence of EXO1. Hence, such a scaffolding function might contribute to the intermediate phenotype we observed in *Exo1^DA/DA^* mice during the error-prone MMR phase that introduces A:T mutations during SHM in B cells.

Further supporting tier-2 in B cells, we also observed independent contributions from the nuclease and scaffolding functions of EXO1 during CSR. Two variants of error-prone DSBR, namely cNHEJ and microhomology-mediated alt-EJ, are known to contribute during CSR, which can be roughly dissected by studying antibody S-S junctions (38). Our data highlight that *Exo1^DA/DA^* mice show a clear preference for blunt-end NHEJ with an average of 0 bp S-S overlap compared to *Exo1^−/−^* B cells that show about 2-3 bp of S-S microhomology overlap. *Exo1^DA/DA^* cells also have a decreased propensity for long microhomologies at the S-S junctions and at the same time an increased propensity for insertions. This again could be explained by a i) passive dominant-negative effect of the *Exo1^DA/DA^* protein manifesting longer than usual residency period on DSB sites. But it could also be argued that ii) in complex scenarios such as during the competition between NHEJ and alt-EJ at S-S junctions, EXO1 can fulfill a more complex role than the basic nuclease activity primarily required for HR. This complexity could be mostly attributed to high selection for NHEJ and alt-EJ at S-S junctions where EXO1 is not expected to fulfill a catalytic role but rather a scaffolding one. The recruitment of EXO1 by CtIP or DNA2 nucleases which are shown to bind, recruit, and synergize with EXO1 nuclease activity, might contribute to this S-S junction process through an EXO1 scaffolding role.

Finally, our data show that while EXO1 catalytic activity is completely dispensible for apparently normal meiotic progression, it is critically dependent on the presence of EXO1 protein. This tier-3 scaffolding EXO1 function is surprising because meiosis is considered a specialized and unique form of a hybrid between error-prone HR and MMR which contributes to organismal diversity (54, 55). It is not clear why EXO1 nuclease activity is not required in meiosis, but it is consistent with budding yeast studies, which confirm our findings and further suggest that the nickase activity of the MLH1-MLH3 complex is responsible for crossover formation; albeit dependent on EXO1 protein but not on EXO1 nuclease activity (29). This could explain why both sexes of *Exo1^−/−^* mice are sterile (56). Furthermore, recent studies in *Exo1^DA/DA^* mice showed that EXO1 is not an important contributor to 5⍰ to 3⍰ end resection in murine meiosis or is substantially redundant with other resection activities (56). In fact, recent studies suggest that EXO1 helps facilitate the MLH1-MLH3 nuclease activity in meiosis (30, 57).

In conclusion, we show that EXO1 fulfills 3-tier activity in genome maintenance depending on the process it is involved in and the complexity of that DNA repair process. One tier is predominantly dependent on the nuclease function of EXO1 such as canonical MMR and DSBR. Another tier depends primarily on the scaffolding function, such as meiosis, and a final tier depends on the concerted act of nuclease and scaffolding functions during non-canonical repair processes in B cells for antibody diversification.

## ACKNOWLEDGMENTS

This work was supported by the NIH Grants CA72649, CA102705 (to MDS) and 1R01AI132507-01A1 (to MDS and TM), CA76329, CA222358 and CA248536 (to WE), and grants from UZH-URPP, BBSRC [BB/N017773/2], and SNF [CRSK-3_190550] (to RC), and grants from Agencia Estatal de Investigación [PID2020-112994RB-I00/AEI/10.13039/501100011033] and Ministerio de Economía, Industria y Competitividad [SAF2017-82309-R/MINECO/AEI/FEDER, EU] (to SR). MDS is supported by the Harry Eagle Chair, provided by the National Women’s Division of the Albert Einstein College of Medicine. SR was supported by Ministerio de Economía y Competitividad through Programa Ramón y Cajal (RYC-2014-16399/MEC). RC was supported by AMS SBF001\1005. RBM was supported by EPSRC EP/M506527/1.

## SUPPLEMENTARY INFORMATION

**Figure S1**: Generation of *Exo1^DA/DA^* mice. A) The mouse Exon1 DA allele was generated by CRISPR/Cas9 mediated gene editing and carries a one-nucleotide substitution mutation (A to C) at codon 173 in exon 6 of the Exon1 gene to generate the amino acid substitution D173A. B) Mouse cohorts were screened using PCR and Sanger sequencing to identify offspring carrying the desired genetic mutation, as depicted. C) Further western blot analysis using mouse embryonic fibroblasts extracts from various cohorts along with an anti-Exo1 antibody confirms the maintenance and stability of Exo1 protein in wild type and DA mutant cells.

