## Supplementary figures and images for "Role of EXO1 nuclease activity in genome maintenance, the immune response and tumor suppression in *Exo1^D173A^* mice"

### Figure S1

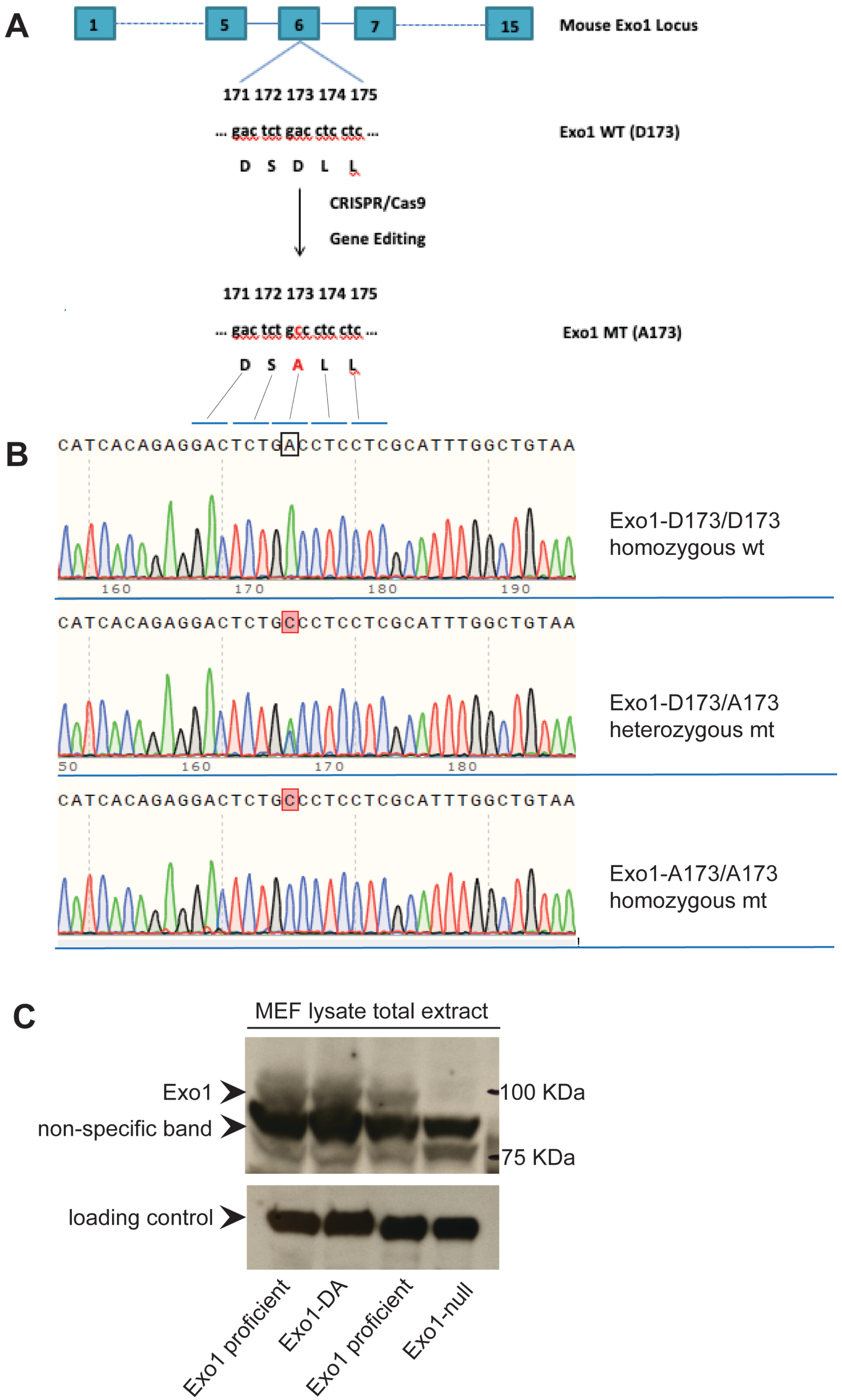
